# Epigraph Hemagglutinin Vaccine Induces Broad Cross-reactive Immunity Against Swine H3 Influenza Virus

**DOI:** 10.1101/2020.05.13.094615

**Authors:** Brianna L. Bullard, Brigette N. Corder, Bette Korber, Eric A. Weaver

## Abstract

Swine influenza virus (SIV) is a significant burden on the pork industry and threat to human health due to its zoonotic potential. Here, we utilized epigraph, a computational algorithm, to design a universal swine H3 (swH3) influenza vaccine. The epigraph hemagglutinin proteins were delivered using an Adenovirus type 5 vector and compared to a wild type hemagglutinin and the commercial inactivated vaccine, FluSure. In mice, epigraph vaccination led to significant cross-reactive antibody and T-cell responses against a diverse panel of swH3 isolates. Epigraph vaccination also reduced weight loss and lung viral titers in mice after challenge with three divergent swH3 viruses. Vaccination studies in swine, the target species for this vaccine, showed stronger levels of cross-reactive antibodies and T-cell responses after immunization with epigraph vaccine compared to the wild type and FluSure vaccines. In both murine and swine models, epigraph vaccination showed superior cross-reactive immunity that should be further investigated as a universal swH3 vaccine.

## Introduction

Influenza infection in swine is a highly contagious respiratory virus endemic in pig populations around the world^1,2^. Swine influenza virus (SIV) can cause zoonotic infections in humans, representing a potential threat to human health^3,4^. When influenza virus of swine origin infects humans, it is termed a variant infection. Since 2010, there have been >460 reported swine influenza virus (SIV) variant infections in humans in the United States of America (USA)^5^. Pigs are susceptible to swine, avian, and human influenza viruses, making them the perfect “mixing vessel” for novel reassorted influenza viruses^3,6^. These novel reassorted viruses have significant pandemic potential if zoonosis occurs, as seen with the 2009 H1N1 “swine flu” pandemic. This highly-reassorted swine-origin influenza virus quickly circulated the globe and infected a staggering 24% of the world’s human population^7,8^. As the first influenza pandemic of the twenty-first century, this highlights the threat that zoonotic SIV poses to human health.

SIV not only poses a potential human health threat from zoonosis, but it also represents a significant burden on the pork industry. SIV infection of pigs results in high morbidity, with many of the same symptoms as human influenza infections^9^. SIV infection can cause tremendous economic loss to swine producers, with cost estimates as high as $10.31 per market pig^10^. In the USA, over 95% of swine nursery sites vaccinated weaned pigs against SIV infection. However, 50% of those sites also reported SIV infections in their herds despite vaccination^11^. This highlights the ongoing challenge of vaccinating against the highly diverse and evolving influenza virus. Currently, most commercial SIV vaccines are traditional whole inactivated virus (WIV) vaccines containing both H1 and H3 subtypes, often with an oil-in-water adjuvant^12^. However, these commercial vaccines are infrequently updated and do not protect against the large diversity of SIV circulating in the swine population. This has led to the use of autogenous, or custom, vaccines which contain herd-specific SIV strains and are limited to use within that herd^13^. An estimated 50% of SIV vaccines sold are autogenous vaccines^11,12,14^. However, autogenous vaccines have multiple drawbacks, including labor intensive laboratory techniques for diagnosis, isolation, virus growth, and purification, which results in a lag period before the vaccine can be administered^12^. The limited strains currently available in commercial swine influenza vaccines paired with the significant drawback to autogenous vaccines highlight the urgent need for a universal swine influenza vaccine. A universal swine influenza vaccine could reduce the economic impact of SIV on the pork industry, along with reducing the risk of emergent zoonotic influenza viruses into the human population.

Currently, the SIV subtypes H1N1, H1N2, and H3N2 circulate in the swine population worldwide^1^. We chose to focus on the swine H3 (swH3) subtype for this study because the H3N2 subtype accounted for >90% of the SIV variant human infections reported in the US since 2010^5^. The swH3 subtype is highly diverse, with contemporary strains circulating in Europe and Asia that are divergent from contemporary strains circulating in North America^15,16^. The North American strains are divided into clusters I-IV, with cluster IV further divided into A-F^15^. The high diversity of the swH3 population represents a significant challenge in the development of a vaccine that induces high-levels of broadly cross-reactive immunity.

This study aims to evaluate a novel vaccine antigen designer, called the Epigraph vaccine designer tool, for the design of a universal swH3 influenza vaccine ^17^. Epigraph is a graph-based algorithm that creates a cocktail of vaccine antigens designed to maximize the potential epitope coverage of a highly diverse population. This epigraph algorithm has been used to predict therapeutic HIV vaccine candidates^18^ and has shown promising potential *in vivo* as a Pan-Filovirus vaccine^19^. Here, we utilize the Epigraph vaccine designer in the development of a universal swH3 vaccine by computationally-designing a cocktail of three swH3 hemagglutinins (HA), a surface glycoprotein of influenza. This is the first report evaluating the epigraph algorithm for the design of a broadly-reactive influenza vaccine. The epigraph HA immunogens were expressed in a replication-defective Adenovirus type 5 (Ad) vector and compared to a wild-type hemagglutinin (HA) expressed in Ad and the commercial inactivated adjuvated vaccine, FluSure. We evaluated the cross-reactivity of the epigraph vaccine by measuring both antibody and T-cells responses in mice and swine. Additionally, we evaluated cross-protective immunity against three diverse swH3 strains after challenge in mice. These data support the use of epigraph immunogens in the development of a universal swH3 vaccine.

## Methods and Materials

### Ethics Statement

Female BALB/c mice ages 6-8 weeks were purchased from Jackson Laboratory. Outbred male and female pigs aged 3 weeks were purchased from Audubon Manning Veterinary Clinic (AMVC). Mice and swine were housed in the Life Sciences Annex building on the University of Nebraska – Lincoln (UNL) campus under the Association for Assessment and Accreditation of Laboratory Animal Care International (AAALAC) guidelines. The protocols were approved by the UNL Institutional Animal Care and Use Committee (IACUC) (Project ID 1217, 1717, and 1879). All animal experiments were carried out according to the provisions of the Animal Welfare Act, PHS Animal Welfare Policy, the principles of the NIH Guide for the Care and Use of Laboratory Animals, and the policies and procedures of UNL.

### Influenza Viruses

The following swine influenza viruses were obtained from the Biodefense and Emerging Infectious Diseases Repository: A/swine/Ohio/09SW73E/2009 (Ohio/2009) [NR-36705], A/swine/Ohio/11SW87/2011 (Ohio/2011) [NR-36715], and A/swine/Manitoba/00446/2005 (Manitoba/2005) [NR-43049]. The following viruses were generous gifts from our collaborators: A/swine/Texas/4199-2/1998 (Texas/1998) strain from Dr. Hiep Vu, A/swine/Colorado/1/1977 (Colorado/1977) strain from Dr. Richard Webby, and the A/swine/Kansas/11-110529/2011 (Kansas/2011) strain from Dr. Wenjun Ma. The following swine influenza viruses were obtained from the USDA Swine Surveillance Influenza A virus isolates repository: A/swine/Minnesota/A01270872/2012 (Minnesota/2012), A/swine/Colorado/23619/1999 (Colorado/1999), A/swine/Wyoming/A01444562/2013 (Wyoming/2013), A/swine/Minnesota/A01432544/2013 (Minnesota/2013), A/swine/Indiana/A01202866/2011 (Indiana/2011), and A/swine/Texas/A01785781/2018 (Texas/2018). The Manitoba/2005, Colorado/1977, and Ohio/2011 swine influenza viruses were mouse-adapted through serial lung passaging in mice 7 times.

The following human influenza viruses were obtained from the Biodefense and Emerging Infectious Diseases Repository: A/Texas/1/77 (Texas/1977) [NR-3604], A/Mississippi/1/85 (Mississippi/1985) [NR-3502], A/Aichi/2/68 (Aichi/1968) [NR-3483], A/Beijing/4/89 (Beijing/1989) [NR-3495], A/Nanchang/933/95 (Nanchang/1995) [NR-3222], A/Ann Arbor/3/93 [NR-3524], and A/Mississippi/1/85 [NR-3502]. The Texas/1977 influenza virus was mouse-adapted through serial lung passaging in mice 5 times. All swine and human viruses were passaged one time in specific pathogen free (SPF) embryonated eggs and the chorioallantoic fluid was stored at −80°C. Viruses were quantified based on HAU and TCID_50_.

### Design and characterization of the Epigraph immunogens

The epigraph vaccine immunogens were designed using the Epigraph Vaccine Designer at the Los Alamos National Laboratories^17,18^. First, all unique swine H3 hemagglutinin (HA) sequences (duplicates excluded) were downloaded from the Influenza Research Database as of April 25^th^, 2017. This resulted in 1,561 HA sequences that where then uploaded to the Epigraph Vaccine Designer and run with the following parameters: epitope length: 9, cocktail size: 3. The resulting cocktail of three HA epigraph genes were added back to the swine H3 sequence population and aligned using ClustalW. Because we included over 1,500 sequences in the analysis, we used a simple Neighbor Joining strategy to build the phylogenetic tree using the Jukes-Cantor model with a Blosum62 cost matrix on the Geneious 11.1.5 software. The cluster designation for each swH3 strain was determined using literature^20^ and location of the phylogenetic tree relative to reference strains. Phylogenetic trees to compare the assay strains to the vaccine strains were created by maximum likelihood estimation using PhyML 3.3 with a Jones-Taylor-Thornton substitution model on the Geneious 11.1.5 software^21^.

### Construction of the replication-defective Adenovirus vectors

The three epigraph HA and wildtype A/swine/Texas/4199-2/1998 HA immunogens were codon-optimized for swine gene expression and synthesized by GenScript. These genes were cloned into an Adenovirus type 5 replication-defective E1/E3 deleted vector using the Ad-Easy Adenoviral Vector System (Agilent). Briefly, the HA genes were cloned in the pShuttle-CMV plasmid and cotransformed with pAd-Easy-1 (Ad5 genome) into BJ5183 cells for homologous recombination into the E1 region of the Ad5 genome. Ad5 recombinants were confirmed by restriction digest and sequencing and then midiprepped using the Qiagen Hi-Speed Midiprep Kit. The recombinant Ad5 genomes with HA inserts were linearized with PacI and buffer exchanged using a Strataprep PCR purification kit (Agilent Technologies). The linearized recombinant gDNA was transfected into 293 cells using the PolyFect Transfection Reagent (Qiagen). After virus rescue was observed via plaque formation, cells were harvested and virus was released by 3 freeze-thaw cycles. Virus was amplified by sequential passages in 293 cells until a final amplification using a Corning 10-cell stack (~6300 cm^2^). The virus was purified by 2 sequential CsCl ultracentrifuge gradients, desalted using Econo-Pac 10DG Desalting Columns (Bio-Rad), and stored at −80°C. Virus particles (vp) were quantitated by OD260. The infectious units per mL was determined using the AdenoX Rapid Titer kit according to the manufacturer’s instructions (Clontech Laboratories).

### Western blot

HA protein expression from the recombinant Ad vectors was confirmed by western blot. Confluent 293 cells were infected at a MOI of 10 and incubated at 37°C and 5%CO_2_ for 48 hrs. Cells were harvested, denatured using Laemmli buffer plus 2-mercaptoethanol, boiled at 100°C for 10 min, and then passed through a QIAshredder (Qiagen). Samples were run on a 12.5% SDS-PAGE gel and transferred to a nitrocellulose membrane. The membrane was blocked for 30 min with 5% milk in TBST before being incubated overnight at 4°C with A/swine/Indiana/0392/2011 polyclonal antibody (BEI NR-50411) at 1:1000 in TBST 1% milk. After 3 washes with TBST, the membrane was incubated for 1 hr at room temperature (RT) with anti-goat HRP conjugated antibody (R&D Systems #HAF109) at 1:2000 in TBST 1% milk. The membrane was washed with TBST and developed with SuperSignal West Pico Chemiluminescent Substrate (Thermo Scientific). A duplicate membrane was run for a GAPDH loading control and probed with anti-GAPDH (Santa Cruz Biotechnology #0411) at 1:1000 and secondary goat anti-mouse HRP conjugated antibody (Millipore Sigma #AP308P) at 1:1000 under the same conditions described above.

### Mouse vaccination and tissue collection

For immune correlate analysis in mice, female BALB/c mice were vaccinated with 10^10^ vp of either Ad-epigraph (the cocktail of three epigraph immunogens at equal ratios to total 10^10^ vp) or Ad-TX98. The commercially available swine influenza vaccine FluSure was administered at 10 times the mouse equivalent dose. All vaccines were compared to a PBS sham vaccinated control group. All immunizations were performed intramuscularly with a 27-gauge needle into both quadriceps in two 25 µl injections. At 3 weeks post-vaccination, mice were bled from the submandibular vein and boosted with the homologous vaccine and dose. Two weeks later, all mice were terminally bled via cardiac puncture and spleens were harvested for analysis of cellular immune response. Sera was isolated from whole blood using a BD Microtainer Blood Collection Tube (Becton Dickinson). To isolated splenocytes, spleens were passed through a 40 μm Nylon cell strainer (BD Labware) and red blood cells were lysed using ACK lysis buffer. Splenocytes were resuspended in cRMPI with 10% FBS and used for ELISpot assays. All mice immunizations and bleeds were performed under isoflurane or ketamine and xylazine induced anesthesia.

### Swine vaccination and tissue collection

For immune correlate analysis in swine, outbred male and female pigs aged 3 weeks were purchased from AMVC. Pigs were pre-screened for swine influenza exposure using an influenza virus nucleoprotein (NP) ELISA and confirmed negative. The pigs were randomly divided into three groups of five and acclimated for 4 days prior to vaccination. Pigs were vaccinated with 10^11^ vp of Ad-epigraph or Ad-TX98 intramuscularly. FluSure animals were vaccinated according to the manufacturer instructions with a 2mL dose intramuscularly. Three weeks later, animals were bled to examine antibody development after a single shot of vaccine and then boosted with the same vaccine and dose as the prime. Sera was isolated from whole blood using BD Vacutainer Serum Separator Tube (Becton Dickinson). Two weeks after boosting, animals were sacrificed to examine humoral and cellular immune correlates. Sera was collected to examine antibody development. In addition, whole blood was collected for isolation of PBMCs using a syringe pre-loading with EDTA. Whole blood was diluted 1:1 with sterile DPBS, gently added on top of lymphocyte separation media (Corning #25072CV) and spun at 400g for 30 min. The PBMC layer was collected, washed with RPMI, and then residual red blood cells lysed with ACK lysis buffer. PBMCs were resuspended in cRMPI with 10% FBS and used for ELISpot assays.

### Hemagglutination inhibition (HI) assay

Sera from mice and swine were incubated with receptor destroying enzyme (RDE; (370013; Denka Seiken) at a 1:3 ratio (sera: RDE) overnight at 37°C followed by inactivation at 56°C for 30 min. Sera was further diluted to a starting ratio of 1:10 with DPBS before use in the HI assay. Sera were serially diluted two-fold in a 96 well V-bottom plate before an equal volume (25 μL) of 4 hemagglutination units (HAU) of virus was added to each well. After incubation at RT for 1 hr, 50 μL of 0.5% chicken red blood cells were added to each well and hemagglutination patterns were read after 30 min.

### ELISpot assay

The T-cell response to vaccination was analyzed using an IFNγ ELISpot assay. Peptide arrays of the HA protein of swine influenza virus strains Ohio/2011, Manitoba/2005, Texas/1998, and Colorado/1977 were synthesized by GenScript. These peptide arrays spanned the entire HA protein of each strain and consist of 17-mers with 10 amino acid overlap. Peptide arrays of the HA protein of human influenza virus strains Texas/1977, Mississippi/1985, and Aichi/1968 were also synthesized by GenScript and were 17-mers with 12 amino acid overlap. Potential immunogenic peptides were identified using a matrix of peptides pools, and the epitopes were confirmed using individual peptides. For ELISpot assays on mice splenocytes, 96-well polyvinylidene difluoride-backed plates (MultiScreen-IP, Millipore) were coated with 50 μl of anti-mouse IFN-γ mAb AN18 (5 µg/ ml; Mabtech) overnight at 4°C before being washed and blocked with cRMPI 10% FBS for 1 hr at 37C. Single-cell suspension of mouse splenocytes were added to each well followed by an equal volume (50 μL) of peptide (5 μg/mL). Plates were incubated overnight at 37°C with 5% CO_2_, washed 6X with PBS, and incubated with 50 μL of biotinylated anti-mouse IFN-γ R4-6A2 mAb (1:1000 dilution; Mabtech) diluted in PBS with 1% FBS for 1 hour at RT. Plates were washed 6X with PBS and incubated with 50 µl of streptavidin-alkaline phosphatase conjugate (1:1000 dilution; Mabtech) diluted in PBS 1% FBS. After 1 hour at RT, the plates were washed 6x with PBS and developed by adding 100 µl of BCIP/NBT (Plus) alkaline phosphatase substrate (Thermo Fisher). Development was stopped by washing several times in dH_2_O. The plates were air dried and spots were counted using an automated ELISpot plate reader (AID iSpot Reader Spectrum). Results are expressed as spot-forming cells (SFC) per 10^6^ splenocytes. Swine ELISpot assays on PBMCs were performed as described above, however, plates were coating with 50 μL of anti-porcine IFN-γ mAb pIFNγ-I (5 µg/ ml; Mabtech) and incubated with 50 μL of biotinylated anti-porcine IFN-γ mAb P2C11 (1:1000 dilution; Mabtech) after overnight incubation. One pig in the epigraph group was excluded from the ELISPOT analysis due to cell viability loss. The MHCI binding predictions were made on 3/5/2020 using the IEDB analysis resource Consensus tool^22^.

### Influenza challenges in mice

BALB/c mice (n=10) were vaccinated with 10^10^ vp of the Ad-epigraph or Ad-TX98 vaccine, the inactivated vaccine FluSure, or with PBS sham vaccine intramuscularly. After 3 weeks, the mice were challenged intranasally with either 10^4^ TCID_50_ of Ohio/2011, 10^5^ TCID_50_ of Manitoba/2005, 10^3.5^ TCID_50_ of Colorado/1977, or 10^4.3^ TCID_50_ of Texas/1977. On day 3 post challenge, five mice from each group were sacrificed and the lungs were collected in order to examine lung viral titers by TCID_50_ and qPCR. The remaining five mice were monitored for weight loss and were euthanized when they lost 25% of their starting weight.

### Tissue culture infectious dose (TCID_50_)

Mouse lungs from day 3 post influenza challenge were homogenized in PBS, centrifuged at 21,000 g for 10 min, and the lung supernatant collected. The lung supernatant sample was diluted 1:10 in a 96 well U bottom tissue culture dish and serially diluted 10-fold before adding 100 μL of 2×10^5^ cells/mL of MDCK cells to each well. The plates were incubated overnight at 37°C with 5% CO_2_ and then washed one time with sterile DPBS before adding DMEM with 0.0002% trypsin to each well. The plates were then incubated another 3 days at 37°C with 5% CO_2_ before adding 50 μL of 0.5% chicken red blood cells to each well and reading the hemagglutination patterns after 30 min.

### qPCR lung viral load quantification

RNA was extracted from day 3 post challenge lung supernatant using the PureLink Viral RNA/DNA Mini Kit according to manufacturer’s instructions (Invitrogen). Real time qPCR was performed using the Luna Universal Probe One-Step RT-qPCR Kit (NEB) run on a QuantStudio 3 Real-Time PCR System (Applied Biosystems) using the following cycling conditions: 55°C for 30 min, 95°C for 2 min, and 40 cycles of 95°C for 15s and 60°C for 30s. Results were compared to a standard curve created using RNA extracted from a known quantity of infectious virus of Manitoba/2005. The universal primer probe set for Influenza A (BEI Resources, NR-15593, NR-15594, NR-15595) was used.

### Microneutralization Titer

Sera was heat inactivated at 56°C for 30 minutes and then 2-fold serially diluted in a sterile 96 well U bottom before the addition of 50 TCID_50_ of virus per well. After 1 hour of incubation at 37°C, 100 μL of MDCK cells (2×10^5^ cells/mL) were added to each well. The plates were incubated overnight at 37°C with 5% CO_2_ and then washed one time with sterile DPBS before adding DMEM with 0.0002% trypsin to each well. The plates were then incubated another 3 days at 37°C with 5% CO_2_ before adding 50 μL of 0.5% chicken red blood cells to each well and reading the hemagglutination patterns after 30 min.

### Statistical Analysis

GraphPad Prism software was used to analyze all data. Data are expressed as the mean with standard error (SEM). HI titers, T-cell data, and lung viral titers were analyzed using one-way ANOVA. A p value < 0.05 was considered statistically significant (*p < 0.05; **p < 0.01; ***p < 0.001; ****p < 0.0001).

## Results

### Development and characterization of the swH3 epigraph HA vaccine

We designed the swine H3 (swH3) epigraph hemagglutinins (HA) using the Epigraph vaccine designer tool, a graph-based algorithm which creates a cocktail of immunogens designed to maximize potential epitope coverage in a population^17,18^. First, the epigraph vaccine designer determines the frequency of each potential epitope of designated length (*k-mer*) in the target population. The algorithm then uses a graph-based approach to trace a path across the HA protein that contains the most common epitopes in the population, resulting in a full-length computationally-designed HA protein (epigraph 1). The first epigraph, by design, tends to be very central in its composition (Fig. 1A). This algorithm then is repeated, to create complementary epigraph sequences that minimize, to the extent possible, potential epitopes contained in the previous epigraph immunogens. In this way, the epigraph 2 and 3 construct generally contain the second and third most common epitopes in the population, respectively. These sequences will appear as outliers in a phylogeny, as their composition reflects different k-mer frequencies from sequences throughout the tree (Fig. 1A). The resulting trivalent set of epigraph sequences provides the optimal coverage of potential linear epitopes in the population for a 3-protein set, minimizes in the inclusion of rare epitopes that might result in type-specific immune responses, and although artificial, each epigraph resembles natural HA proteins to enable both the induction of antibody and T-cell responses.

**Figure 1.**
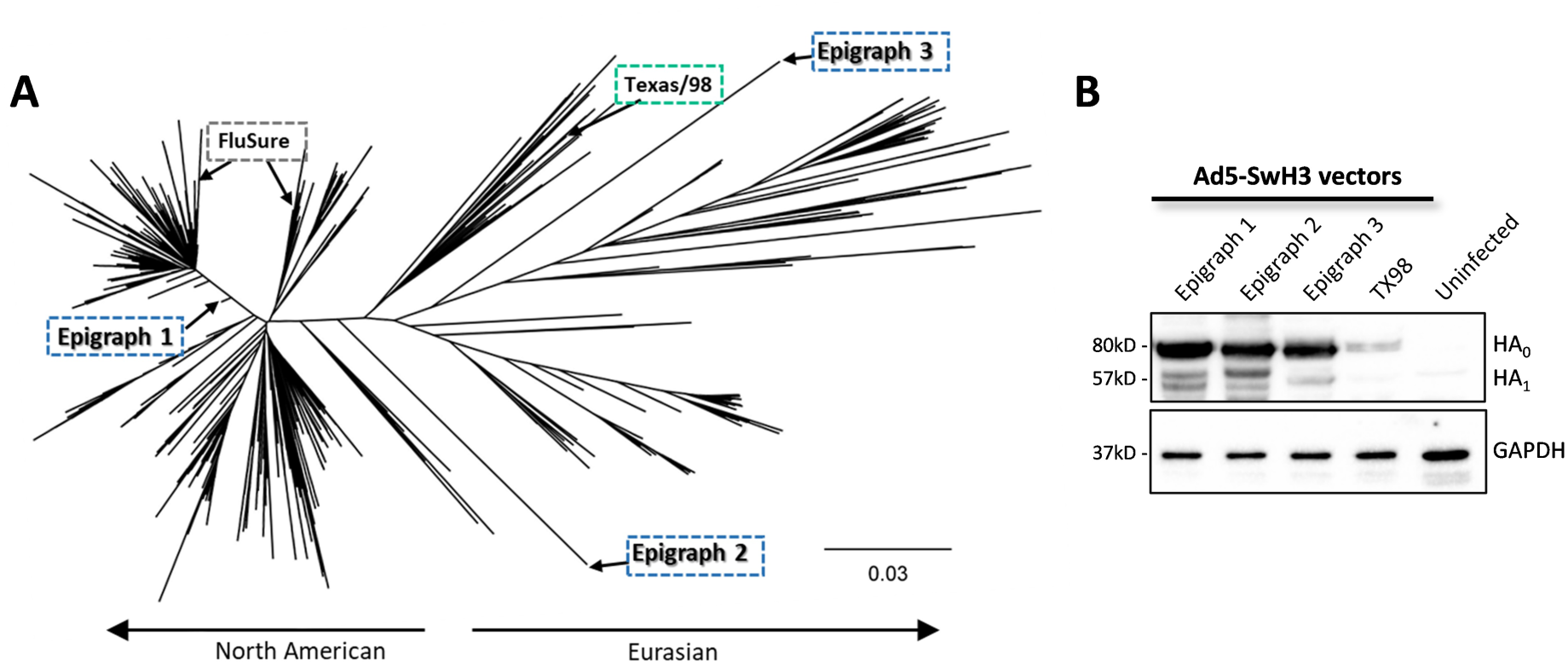
Characterization of the epigraph vaccine constructs. (A) The three swH3 epigraph immunogens were computationally-designed using the Epigraph Vaccine Designer Tool to create a cocktail of immunogens designed to maximize potential epitope coverage in a population. The three epigraph HA immunogens were aligned to the 1,561 unique swine H3 HA sequences using a ClustalW alignment. A neighbor-joining tree was constructed to visualize the phylogenic relationship between the vaccine immunogens and the population of swH3 sequences. The three epigraph immunogens, the Texas/1998 (TX98) wild-type HA comparator, and the two FluSure strains are labeled for reference on the phylogenetic tree. Epigraph, wildtype and FluSure vaccines are shown in the blue, green and black boxes, respectively. (B) All three epigraph immunogens and the TX98 HA were cloned into a replication-defective Adenovirus type 5 (Ad) vector and HA protein expression was confirmed by Western Blot. GAPDH is used as a cellular protein loading control.

The resulting three epigraph HA sequences were aligned back to the original swH3 sequence population and a phylogenic tree was constructed to visualize their relationship to the swH3 population. The three epigraph swH3 immunogens localize across the phylogenic tree (Fig. 1A). In order to evaluate the computational design of the epigraph vaccine, we selected a HA gene which localizes near the center of the tree (A/swine/Texas/4199-2/1998 [TX98]) as a wild type comparator. In addition, we also compared our epigraph vaccine to a commercial SIV vaccine, FluSure. FluSure is an inactivated, oil-in-water adjuvanted vaccine which contains two North American swH3 strains (along with two H1 strains), which belong to the IV-A and IV-B clusters. The three swH3 epigraph genes and the TX98 wild-type HA comparator were cloned into a replication-defective Adenovirus type 5 vector (Ad) for gene expression. Gene expression was confirmed via western blot (Fig. 1B) and virus particle (vp) to infectious unit ratios were determined to confirm approximate infectivity between the stocks (**Supp. Table 1**).

### Vaccination with epigraph lead to the development of a strong cross-reactive antibody response in mice

We first evaluated the immune response after vaccination in mice. BALB/c mice (n=10) were vaccinated with 10^10^ virus particles (vp) of the Ad-epigraph vaccine, which consisted of equal ratios of the three Ad-epigraph viruses totaling 10^10^ vp. Our epigraph vaccine was compared to mice vaccinated with either 10^10^ vp of the Ad-TX98 wild type comparator or 50 uL of FluSure (which translates to 10X the equivalent dose of a three week old pig). A PBS sham vaccine was used as a negative control. Three weeks later, mice were boosted with the same vaccine. Mice were sacrifice two weeks after boosting to examine the humoral and cellular immune response after vaccination (Fig. 2A). The cross-reactivity of the antibody response was examined using an hemagglutination inhibition (HI) assay. We selected a panel of 12 swH3 strains which represented much of the diversity of the swH3 phylogenetic tree, spanned multiple clusters, and included both contemporary and historical isolates. A phylogenetic tree was constructed to examine the relationship of the selected 12 strains to the vaccine strains (Fig. 2B; **Supp. Table 2**). Vaccination with the epigraph immunogens resulted in a strong cross-reactive antibody response, with HI titers ≥40 to 11 of the 12 (92%) swH3 strains and significantly higher antibody titers compared to the TX98 and FluSure groups for 9 of the 12 of the swH3 strains (Fig. 2C). In contrast, the TX98 wild type comparator and FluSure vaccinated mice developed strong antibody titers (≥40) to 3 of the 12 (25%) swH3 strains. The TX98 group developed a strong antibody response to the matched virus Texas/1998 and limited cross-reactivity with only two other strains (Wyoming/2013 and Minnesota/2012). The FluSure vaccine group developed a strong antibody response to both cluster IV-A viruses and to the Minnesota/2012 cluster IV-B strain (a match for the vaccine strain). However, FluSure vaccination provided only limited cross-reactivity with mismatched viruses.

**Figure 2.**
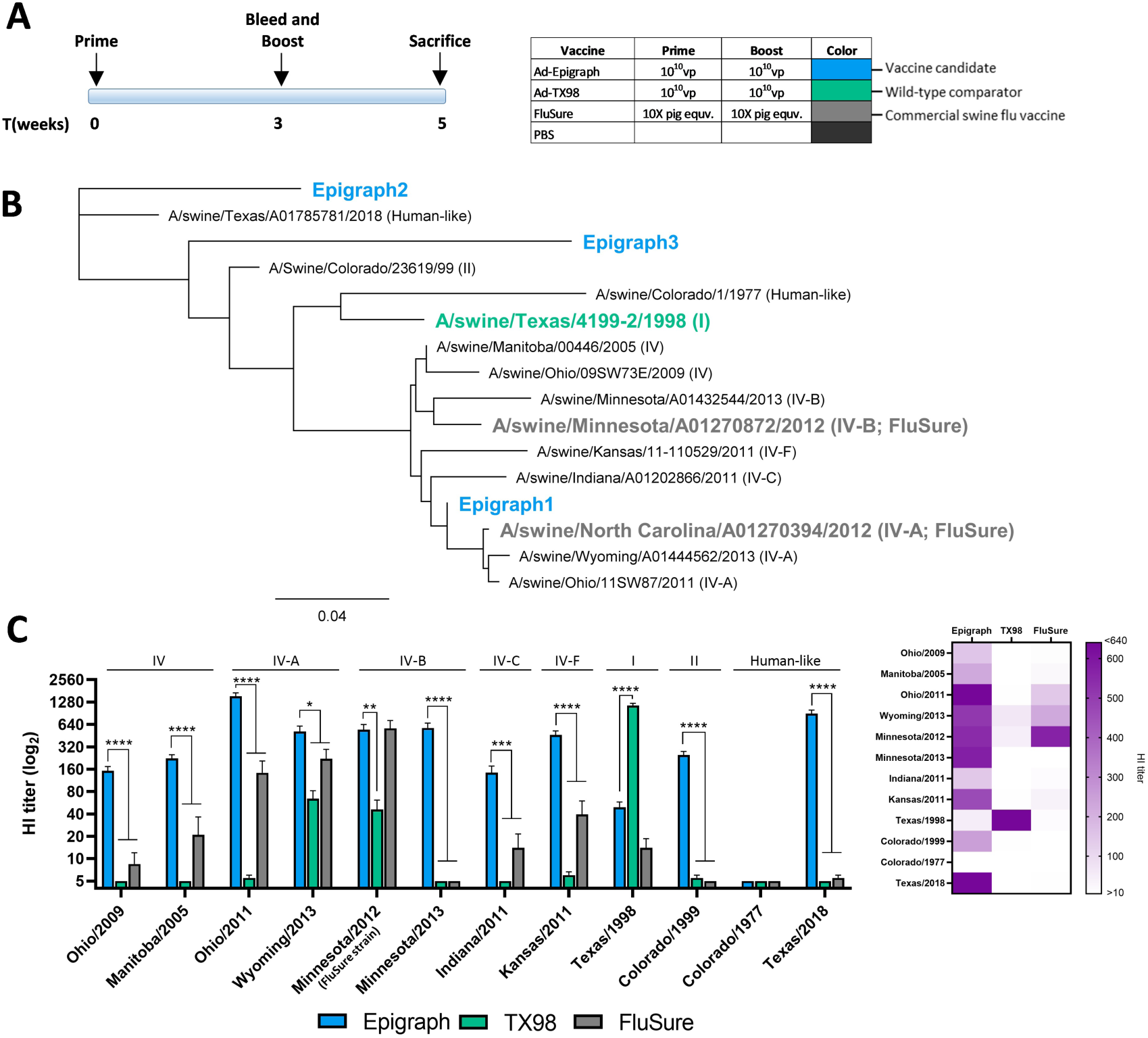
Cross-reactive antibody responses with swH3 strains after vaccination in mice. (A) BALB/c mice (n=10) were vaccinated according to the timeline and vaccine dose. (B) To examine the cross-reactivity of the antibody response after vaccination, a panel of 12 swH3 strains were selected that span the phylogenic tree and represent multiple clusters. A maximum likelihood tree was constructed to visualize the relationship between these assay strains and the vaccine immunogens. The cluster designation is in parentheses after the full strain name. (C) Two weeks after boosting, mice were sacrificed and sera was analyzed using a HI assay against the 12 swH3 representative strains. Cluster designation can be seen above the HI titer bars for each strain. A heat map of these HI titers was constructed to further visualize the total cross-reactive antibody response of each vaccine (one-way ANOVA; *p < 0.05; **p < 0.01, ***p < 0.001, ****p < 0.0001).

### Epigraph immunized mice have a higher total T cell response and recognize more epitopes from four divergent swH3 strains

Cross-reactive T-cells have been shown to play an important role in viral clearance during influenza virus infection^23,24^. Therefore, we wanted to evaluate if there was increased cross-reactivity of T-cell responses after vaccination with the epigraph vaccine. To examine the cross-reactivity, we selected four swH3 strains that represent a large portion of swH3 diversity. Peptide arrays for the Ohio/2011 strain (cluster IV-A), Manitoba/2005 (cluster IV), Texas/1998 (cluster I), and Colorado/1977 (human-like) were constructed. The T-cell response to each of the four strains was mapped using an IFNγ ELISPOT with an overlapping peptide array containing 17-mers with 10 amino acid overlap. Peptides were considered positive if the response was greater than 50 spot forming cells (SFC) per million. Epigraph vaccinated mice recognized a greater number of epitopes across all four swH3 strains as compared to the TX98 vaccinated mice (Fig. 3A). Ironically, the Colorado/1977 strain was the only strain that the epigraph vaccine did not show cross-reactive antibody responses against (Fig. 2C). However, here we see that there is a significant and robust T cell response to the Colorado/1977 virus. Therefore, this strain was selected specifically to examine the potential for cross-reactive T-cells in the absence of detectable cross-reactive antibodies. FluSure vaccinated mice did not develop significant T-cell responses after vaccination. The magnitude of the responses to each peptide revealed an immunodominant epitope in the HA1 region (amino acid 120-128 of the HA protein) that was positive in all four strains after vaccination with epigraph. This epitope was predicted to bind strongly to the MHC-I complex of BALB/c mice22,25 and, therefore, is likely an immunodominant CD8 epitope (Fig. 3B; **Supp. Fig. 1**).

**Figure 3.**
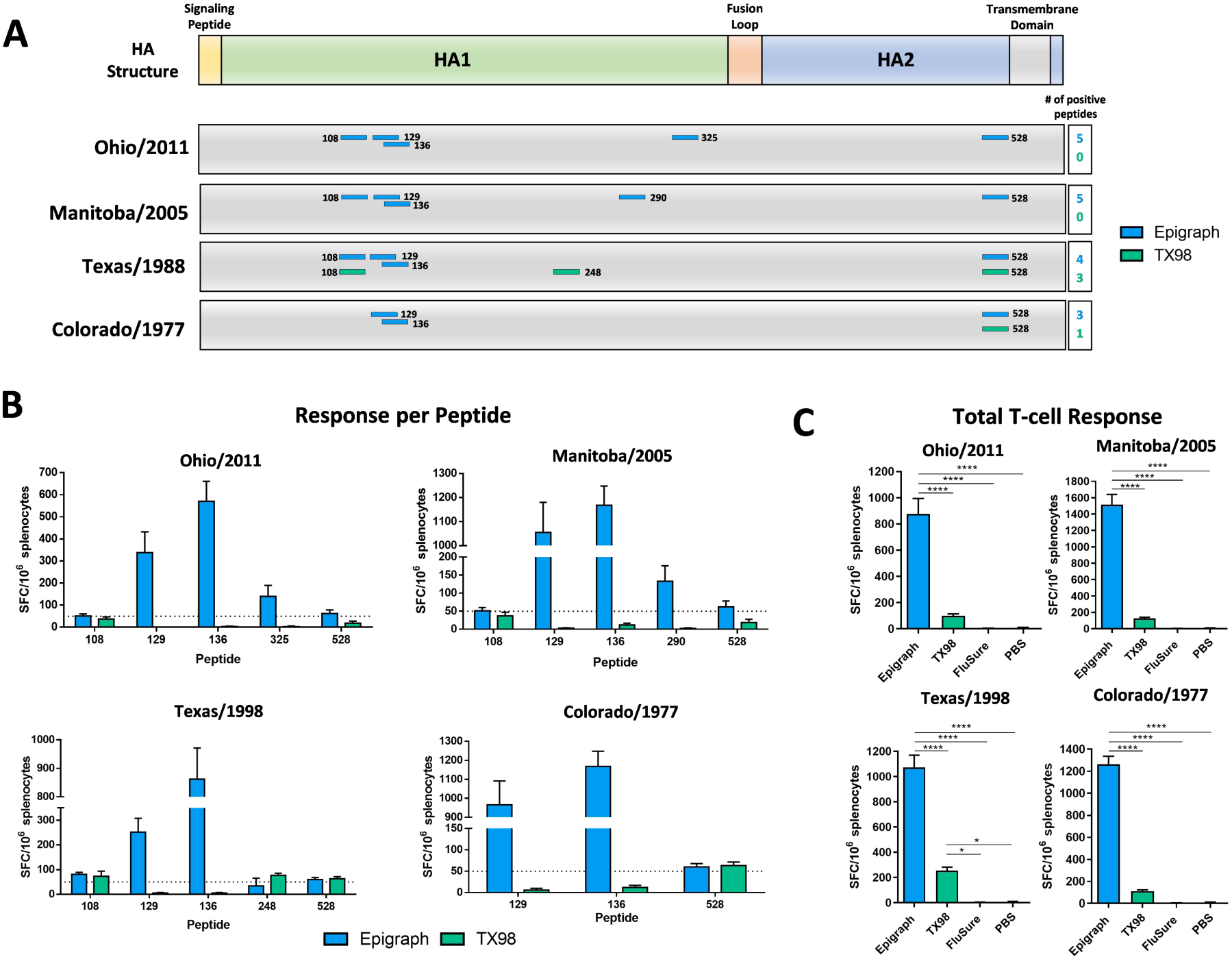
T-cell epitope mapping of four diverse swH3 strains after vaccination. Splenocytes from vaccinated BALB/c mice (n=5) were isolated and analyzed for cellular immunity using an IFNγ ELISpot. T-cell epitopes against the Ohio/2011, Manitoba/2005, Texas/1998, and Colorado/1977 strains were mapped using an overlapping peptide array consisting of 17-mers with 10 amino acid overlap which spanned the entire HA protein. Peptide responses >50 spot forming cells (SFC) per million were considered positive. (A) Positive peptides for each vaccine and their location on the HA protein are indicated. The peptide number designates the position of the last amino acid in the peptide on the total HA protein. (B) The level of response against each positive peptide are reported as SFC per million splenocytes with the dotted line indicating the 50 SFC/million cutoff. (C) The total T-cell response to each virus peptide array is shown for all vaccination groups (one-way ANOVA; *p < 0.05; **p < 0.01, ***p < 0.001, ****p < 0.0001).

Interestingly, T-cells from epigraph vaccinated mice recognized this immunodominant epitope in the Texas/1998 peptide array, however, T-cells from TX98 vaccinated mice did not. One possible explanation may be differences in peptide processing and presentation which are dependent on surrounding sequences. Overall, the total T-cell response was significantly stronger in epigraph vaccinated mice against all four swH3 strains (Fig. 3C).

### Vaccination with epigraph reduces weight loss and lung viral titers after swH3 challenge in mice

We next wanted to determine if the increased cross-reactive antibody and T-cell responses translated to increased protection from a panel of diverse swH3 strains. BALB/c mice (n=10) were vaccinated with a single shot of 10^10^ vp of Ad-epigraph or Ad-TX98, FluSure, or a PBS sham vaccine. Mice were then challenged three weeks later with the mouse-adapted swH3 challenge viruses (Fig. 4A). To examine the antibody response after a single immunization, sera at the time of challenge was examined using an HI assay against each of the three challenge strains (Fig. 4B). A single immunization of epigraph resulted in strong HI titers ≥40 to both Ohio/2011 and Manitoba/2005. In contrast, TX98 vaccination did not result in any detectable antibody responses to these 3 viruses, while FluSure vaccination resulted in low titers (≥40) to Ohio/2011 and Manitoba/2005. No vaccine groups developed antibody responses to the Colorado/1977 strain, making this an ideal strain to evaluate the potential contribution of cross-reactive T-cell responses. After challenge, mice were monitored for weight loss over two weeks. On day three post-challenge, five mice were sacrificed to examine lung viral titers. We measured lung viral titers by both TCID_50_ and qPCR in order to evaluate infectious virus and viral RNA copies, respectively (Fig. 4).

**Figure 4.**
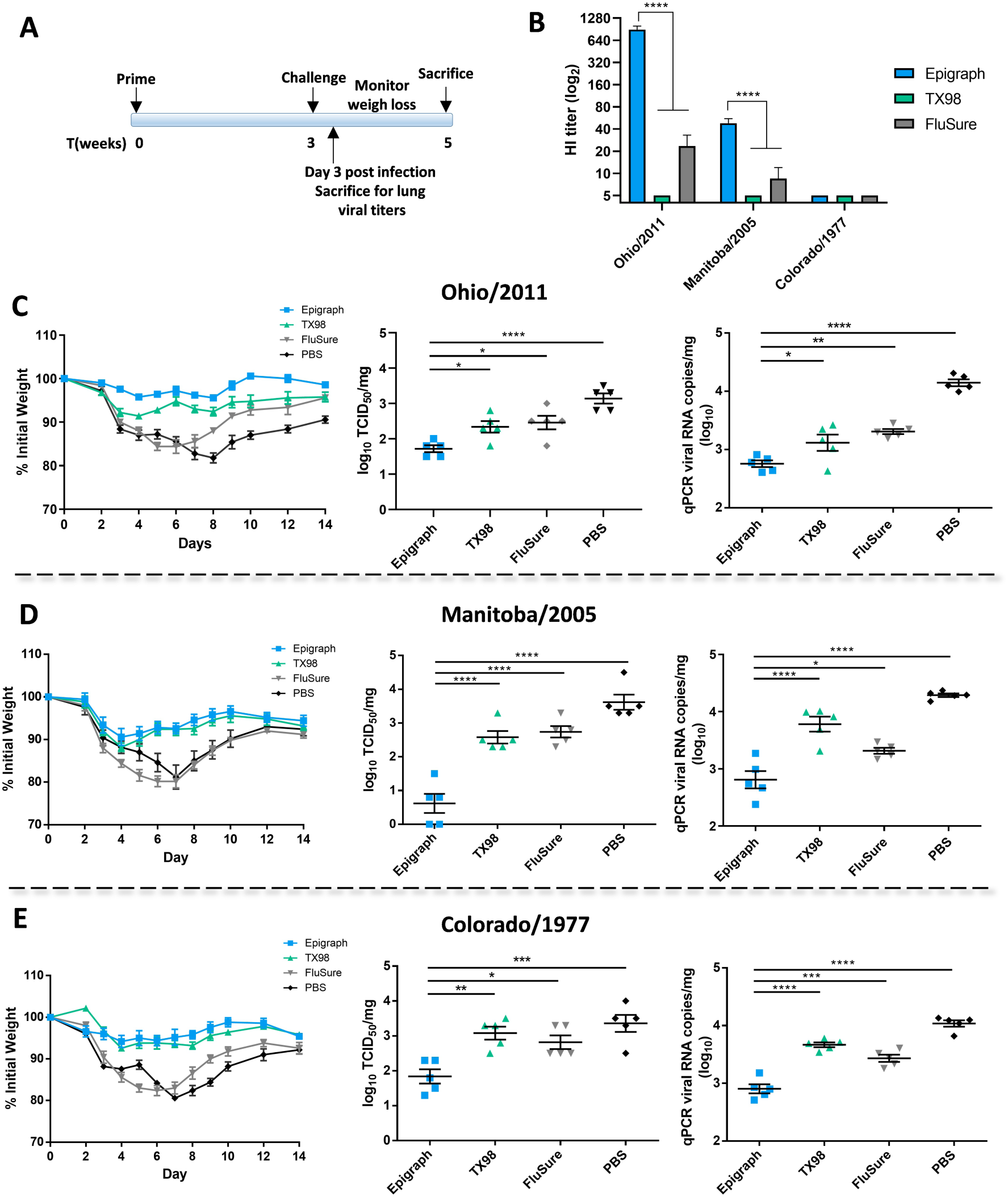
Protection against challenge with divergent swH3 viruses. (A) BALB/c mice (n=10) were vaccinated with 10^10^ vp of Ad-epigraph or Ad-TX98, the commercial inactivated FluSure, or a PBS sham vaccine and challenged according to the timeline (B) An HI titer on mice sera (n=10) was performed to examine the antibody response to the challenge virus strains after a single immunization. Mice were challenged intranasally with 10^4^ TCID_50_ of Ohio/2011 (C), 10^5^ TCID_50_ of Manitoba/2005 (D), or 10^3.5^ TCID_50_ of Colorado/1977 (E) and monitored for weight loss over 14 days. Mice that reached 25% weight loss were humanely euthanized. Three days post infection, five mice per group were sacrificed to examine lung viral titers by TCID_50_ and qPCR (one-way ANOVA compared to the epigraph group; *p < 0.05; **p < 0.01, ***p < 0.001, ****p < 0.0001).

After challenge with the Ohio/2011 strain (cluster IV-A), only epigraph vaccination completely protected mice from weight loss (Fig. 4C). In contrast, the TX98, FluSure, and PBS vaccinated mice lost 8-12% of their body weight by day 3. The FluSure vaccine contains a similar cluster IV-A strain and, although mice were not protected from initial weight loss, the mice showed faster recovery by day 8 as compared to the PBS vaccinated mice (p < 0.0001). In addition, epigraph vaccinated mice showed significantly reduced day 3 lung viral titers as compared the TX98, FluSure, and PBS vaccinated mice (Fig. 4C).

Challenge with the Manitoba/2005 strain (cluster IV) resulted in severe weight loss for the FluSure and PBS vaccinated mice, whereas epigraph and TX98 vaccinated mice were protected from weight loss (Fig. 4D). However, epigraph vaccinated mice showed the lowest lung viral titers on day 3 post challenge as compared to the three other vaccine groups. Interestingly, although both epigraph and TX98 vaccination protected from weight loss, there were significantly higher lung viral titers in the TX98 vaccinated group, supporting that pathology does not always correlate with lung viral titer^26,27^.

Lastly, we challenged with the highly divergent Colorado/1977 strain. All vaccination groups lost weight early after challenge, however epigraph and TX98 vaccinated mice showed significantly reduced weight loss by day 6 as compared to the FluSure and PBS vaccinated mice (Fig. 4E; p < 0.001). Since epigraph and TX98 vaccination does not induce detectable anti-Colorado/1977 antibody responses, the early weight loss but increased recovery could be a result of T-cell mediated protection. Again, epigraph vaccinated mice showed significantly lower lung viral titers on day 3 as compared to the TX98, FluSure, and PBS vaccination mice (Fig. 4E).

### Epigraph vaccination leads to cross-reactive antibody and T cell responses against multiple human H3 strains

Reverse zoonosis, the transmission of influenza virus from human-to-swine, is a key factor in driving the diversity of SIV in swine ^2,28,29^. Therefore, we wanted to determine if our swH3 epigraph vaccine might induce cross-reactive immune responses to human H3 (huH3) isolates in order to reduce reverse zoonotic events. We selected a panel of 7 huH3 strains to evaluate cross-reactive antibody responses by HI assay. A phylogenetic tree was constructed to examine the relationship of these 7 strains to the vaccine strains (Fig. 5A). Epigraph vaccination lead to strong antibody titers ≥40 to 3 of the 7 (43%) huH3 strains (Fig. 5B). TX98 vaccination resulted in antibody titers ≥40 to 2 of the 7 (29%) strains, and is closely related to both strains (>96% identity; Fig. 5A, **Supp. Table 3**). In contrast, FluSure vaccination did not result in cross-reactive antibody responses to any of the huH3 isolates.

**Figure 5.**
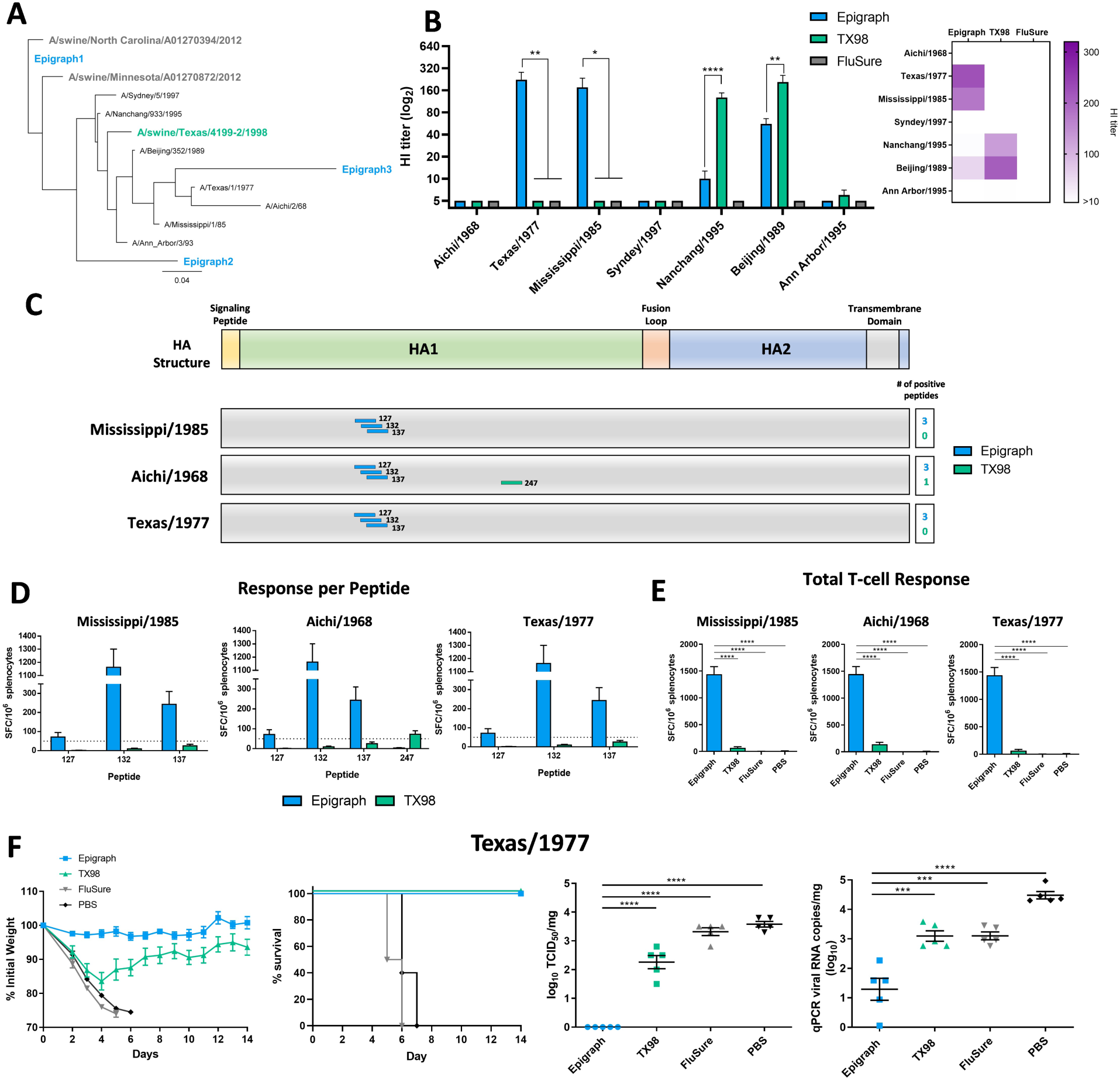
Cross-reactive immune correlates and protection to human H3 isolates. (A) To determine the cross-reactive immune responses of the swH3 vaccines to huH3 isolates, a panel of 7 representative huH3 strains were selected. A maximum likelihood phylogenetic tree was constructed to visualize the relationship of these huH3 assay strains with the swH3 vaccine immunogens. (B) An HI titer was performed against these huH3 strains with sera from BALB/c mice (n=5) vaccinated in fig. 2. A heat map of these HI titers was constructed to further visualize the total cross-reactive antibody response of each vaccine. Splenocytes isolated from vaccinated mice were also examined for cross-reactive cellular immune responses to three huH3 strains using an IFNγ ELISpot. T-cell epitopes against the Mississippi/1985, Aichi/1986, and Texas/1977 strains were mapped using an overlapping peptide array. Peptide responses >50 spot forming cells (SFC) per million were considered positive. (C) Positive peptides for each vaccine and their location on the HA protein are indicated. The peptide number designates the position of the last amino acid in the peptide on the total HA protein. (D) The level of response seen against each positive peptide are reported as SFC per million splenocyte with the dotted line indicating the 50 SFC/million cutoff. (E) The total T-cell response to each virus peptide array is shown for all vaccination groups. (F) BALB/c mice (n=10) were vaccinated with 10^10^ vp of Ad-epigraph or Ad-TX98, the commercial inactivated vaccine FluSure, or a sham PBS vaccine and then challenged three weeks later with 10^4.3^ TCID_50_ of Texas/1977. Mice were monitored for weight loss and sacrificed humanely when 25% weight loss was reached. Five mice per group were sacrificed on day 3 post infection to examine lung viral titer by TCID_50_ and qPCR (one-way ANOVA compared to the epigraph group; *p < 0.05; **p < 0.01, ***p < 0.001, ****p < 0.0001).

We also evaluated cross-reactive T-cell responses to three huH3 strains (Mississippi/1985, Aichi/1968, and Texas/1977) using an IFNγ ELISPOT with overlapping peptide arrays. T-cell mapping was performed as described with the swH3 isolates. Interestingly, epigraph vaccination induced a T-cell response against a single immunodominant epitope present in all three huH3 isolates (Fig. 5C, D). This epitope is the same position as the immunodominant epitope induced against the swH3 isolates (amino acid position 120-128). In the huH3 population, the amino acids in this epitope are highly conserved (~94%; **Supp. Fig. 2**). TX98 vaccination did not induce a T-cell response against this immunodominant epitope and FluSure vaccination did not result in a significant T-cell response against any of the huH3 strains. Epigraph vaccination also resulted in a highly significant total T-cell responses against all three huH3 isolates (Fig. 5E).

To determine if the huH3 cross-reactive immune responses resulted in protection, we challenged vaccinated mice with a mouse-adapted huH3 Texas/1977 isolate. BALB/c mice (n=10) were vaccinated with a single shot of 10^10^ vp of Ad-epigraph or Ad-TX98, FluSure, or a PBS sham vaccine. Mice were then challenged three weeks later with the Texas/1977 challenge strain. Only epigraph vaccination completely protected mice from weight loss and death (Fig. 5F). In contrast, TX98 vaccinated mice lost >16% of their starting body weight before starting to recover. FluSure and PBS vaccinated mice quickly lost weight and were all humanely euthanized by day 7 post infection. Epigraph vaccination also reduced infectious virus in the lungs to undetectable levels on day 4 as measured by TCID_50_ (Fig. 5F).

### Epigraph vaccination in swine induced strong cross-reactive antibody and T-cell responses

Lastly, to confirm that the results seen in mice translated to the target animal, we vaccinated three-week-old pigs intramuscularly with 10^11^ vp of our Ad-epigraph vaccine and compared the immune responses to swine vaccinated with 10^11^ vp of the Ad-TX98 wild type comparator or the commercial vaccine FluSure at the manufactures recommended dose. Three week later, serum was collected to examine the antibody response after a single immunization (Fig. 6A). A single immunization of the epigraph vaccine lead to strong cross-reactive antibody titers ≥40 to 9 out of 12 (75%) of the swH3 strains, with significantly higher antibody responses as compared to the TX98 and FluSure groups to 8 out of 12 of the swH3 strains tested. In contrast, TX98 only resulted in strong antibody titers (≥40) to the matched Texas/1998 strain and FluSure vaccination did not result in significant titers to any of the swH3 after a single immunization.

**Figure 6.**
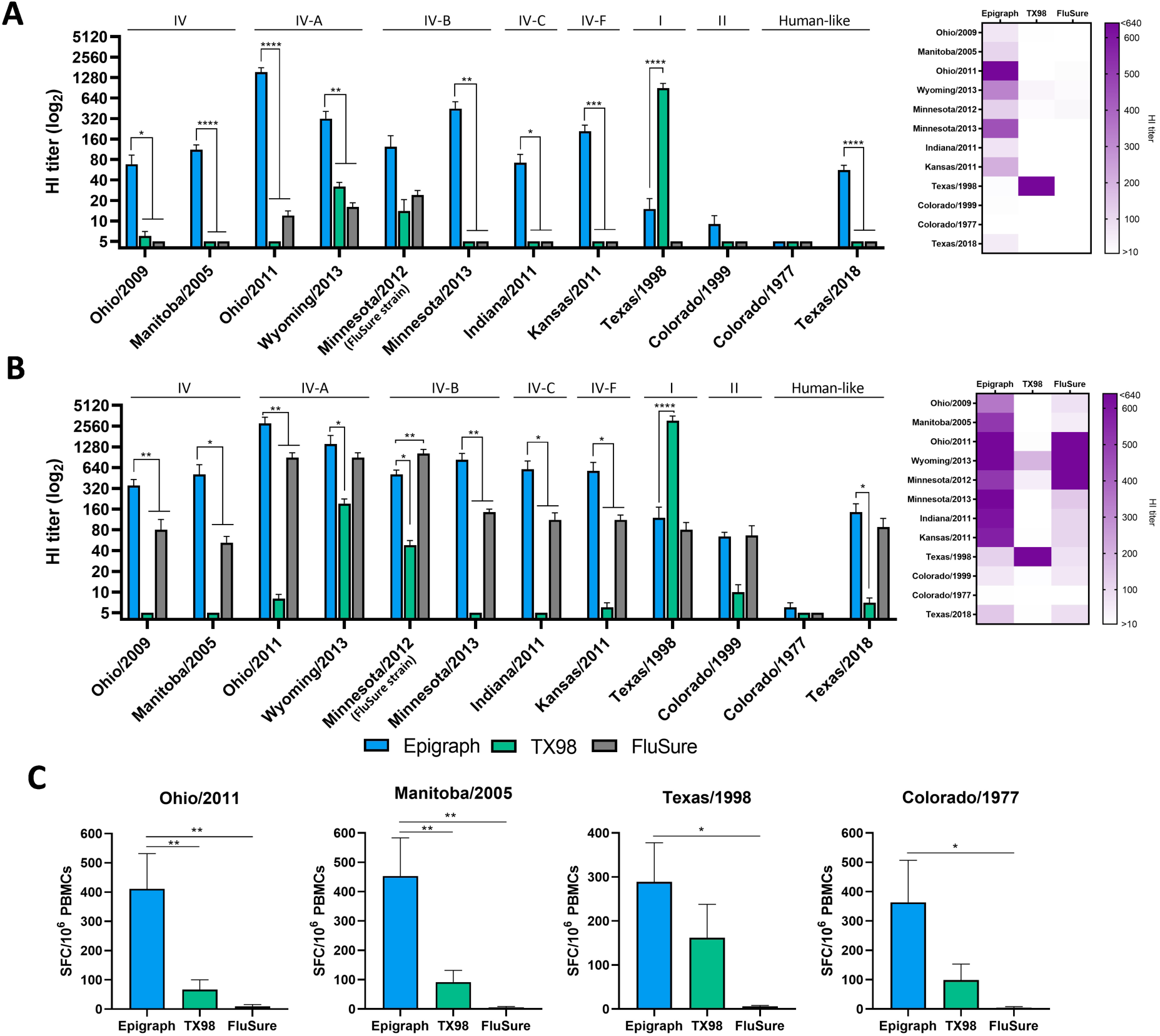
Immune responses to swH3 strains after vaccination in swine. To confirm that the cross-reactive immune responses observed after epigraph vaccination in mice translated to the target animal, three-week-old swine (n=5) were vaccinated with 10^11^ vp of Ad-epigraph or Ad-TX98 or the commercial inactivated vaccine FluSure according to the manufacturer’s instructions. Pigs were bled three weeks later to examine the antibody response after a single shot and then boosted with the same vaccine and dose. Two weeks after boosting, swine were humanely sacrificed. Sera from the single shot (A) or after boosting (B) was analyzed using an HI assay against the 12 swH3 representative strains. Cluster designation can be seen above the HI titer bars. A heat map of these HI titers was constructed to further visualize the total cross-reactive antibody response of each vaccine. (C) PBMCs were isolated to determine the total T-cell response against the four representative swH3 strains (Ohio/2011, Manitoba/2005, Texas/1998, and Colorado/1977) using an IFNγ ELISpot (epigraph n=4; TX98 and FluSure n=5; one-way ANOVA compared to the epigraph group; *p < 0.05; **p < 0.01, ***p < 0.001, ****p < 0.0001).

Pigs were boosted with the same vaccine and dose three weeks after priming and sacrificed two weeks later to examine immune correlates at peak immunity. A second immunization boosted cross-reactive antibody titers in epigraph vaccinated pigs, with titers ≥40 to 11 of the 12 (92%) strains (Fig. 6B). In addition, epigraph vaccination showed significantly higher antibody titers to 9 of the 12 strains as compared to TX98 vaccination and significantly higher antibody titers to 6 of the 12 strains as compared to FluSure vaccination. In contrast, after boosting, the TX98 vaccinated pigs showed strong antibody titers (≥40) to 3 of the 12 (25%) swH3 strains. Interestingly, the TX98 vaccine boost displayed increased antibody titers to the matched Texas/1998 strain along with increased cross-reactivity with the Wyoming/2013 and Minnesota/2012 viruses, the same viruses exhibiting cross-reactivity in the mouse model. The strongest antibody responses induced after boosting with FluSure were against similar strains to the vaccine, the cluster IV-A viruses (Ohio/2011 and Wyoming/2013) and the matched FluSure virus (Minnesota/2012; cluster IV-B). Interestingly, boosting with FluSure also increased the cross-reactive antibody responses across the swH3 panel, with titers ≥40 to 11 of the 12 (92%) strains. However, the responses to most unmatched viruses were significantly lower than responses to epigraph immunization, with an average of 4-fold reduction in HI titers. Indeed, a single immunization of Ad-epigraph resulted in comparable cross-reactive antibody levels as two FluSure immunizations.

To confirm that the cross-reactive antibody responses as measured by HI assay were also functionally neutralizing, we performed a microneutralization assay with the panel of 12 swH3 viruses. Neutralization titer patterns matched those seen in the HI assay, confirming the functionality of these cross-reactive antibodies (**Supp. Fig. 4**). PMBCs were also collected two weeks after boosting to examine the cellular immune response using an IFNγ ELISpot. Epigraph vaccination induced the strongest total T-cell response to all four swH3 strains tested (Fig. 6C). TX98 vaccination resulted a strong T-cell response against the matched Texas/1998 strain but only modest cross-reactive T-cell levels to the other three swH3 strains. FluSure vaccination did not result in detectable cross-reactive T-cell responses.

The post-vaccination swine sera were also examined for the presence of cross-reactive antibodies to the panel of 7 huH3 isolates. After a single immunization, epigraph resulted in strong cross-reactive antibody titers ≥40 to 3 of the 7 (43%) huH3 viruses, the same viruses exhibiting cross-reactivity in the mouse model (**Fig. 7A**). In contrast, TX98 vaccination resulted in antibody titers ≥40 to 1 of the 7 (14%) huH3 strains and FluSure vaccination did not show any cross-reactive antibodies to the huH3 isolates after a single immunization. After a second immunization, the cross-reactive antibody levels in all three vaccine groups increased. Boosting with epigraph resulted in strong antibody titers ≥40 to 6 of the 7 (86%) of the huH3 viruses, while TX98 and FluSure boosting results in antibody titers ≥40 to 3 of the 7 (43%) and 4 of the 7 (57%) huH3 viruses, respectively. However, epigraph showed significantly higher antibody titers to 3 of the 7 isolates as compared to both TX98 and FluSure vaccination.

**Figure 7.**
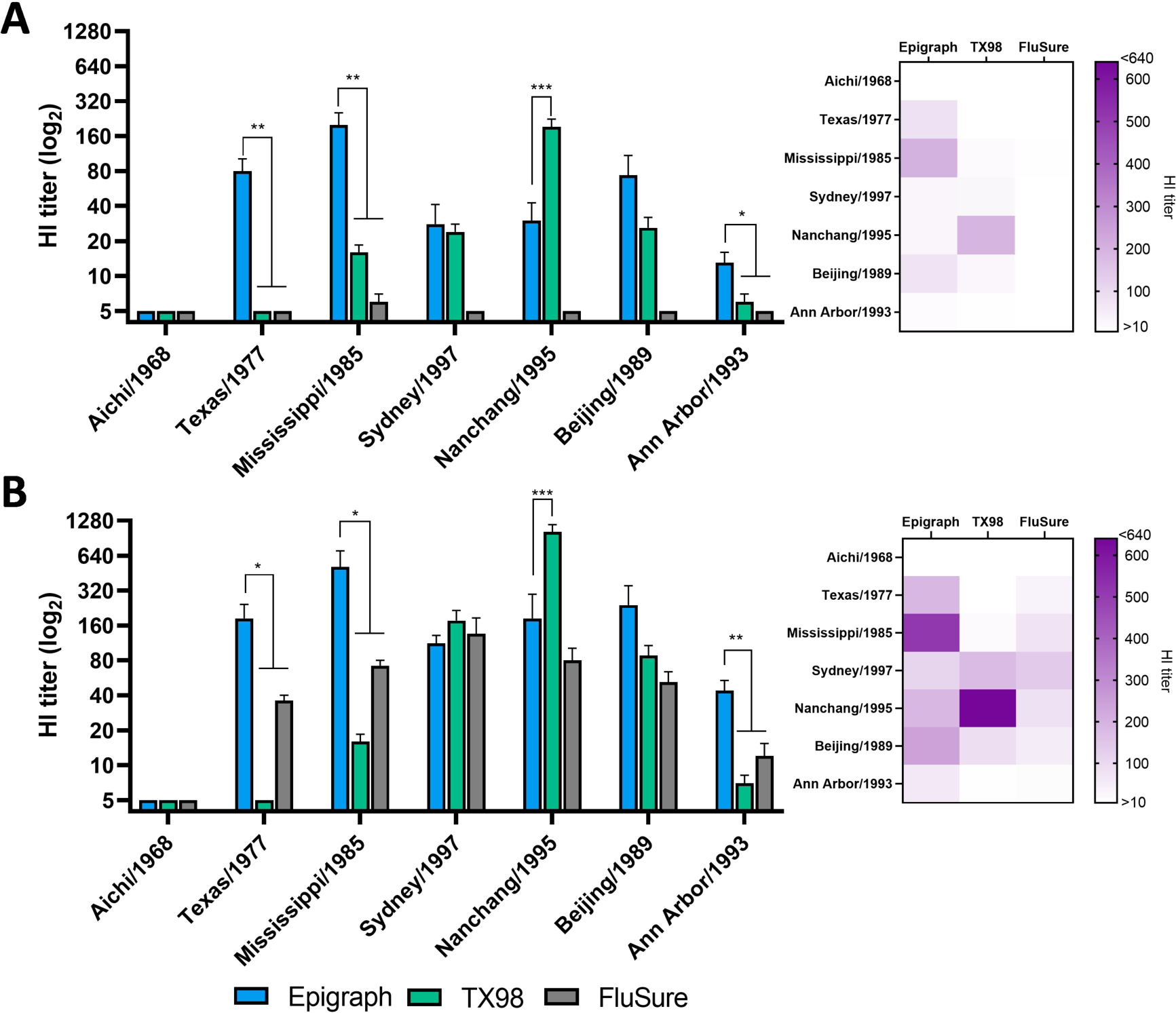
Immune responses to human H3 strains after vaccination in swine. The sera from the vaccinated swine (n=5) was analyzed for cross-reactivity to huH3 strains using an HI assay against the panel of representative 7 huH3 strains. Antibody responses were examined after a single shot (A) or boosting (B) and a heat map of these HI titers was constructed to further visualize the total cross-reactive antibody response of each vaccine (one-way ANOVA compared to the epigraph group; *p < 0.05; **p < 0.01, ***p < 0.001, ****p < 0.0001).

## Discussion

In this study, we evaluated the epigraph vaccine designer algorithm for the immunogen design of broadly cross-reactive swH3 hemagglutinins to create a universal swH3 vaccine. The ideal SIV vaccine would induce protective immunity after a single immunization while also providing broad protection against ever evolving field strains^30,31^. Here, we demonstrated that our epigraph vaccine induced strong cross-reactive antibody responses to a panel of diverse swH3 viruses which represented a large portion of the swH3 diversity. After a prime-boost immunization in mice, the epigraph vaccine induced strong cross-reactive antibody titers to 11 of the 12 highly divergent swH3 viruses. In contrast, the TX98 and FluSure vaccine showed limited cross-reactivity outside of the strains contained in each vaccine. The greater cross-reactive immunity induced by epigraph vaccination in mice was further supported by immunization studies in swine. Importantly, a single immunization with epigraph resulted in strong cross-reactive antibody titers whereas vaccination with TX98 or FluSure showed limited antibody development against unmatched strains. This data suggests that the Ad-epigraph vaccine could be implemented as broadly cross-reactive vaccine which requires only a single immunization for induction of strong immunity.

Interestingly, after the second immunization in swine, the FluSure vaccine showed an increase in breadth of antibody responses. However, the magnitude of the responses against unmatched strains were on average 4-fold lower compared to boosting with the epigraph vaccine. The FluSure vaccine contains two strains of swH3 and an oil-in-water adjuvant which could contribute to the increased antibody breadth observed after boosting^26,32^. Importantly, this data demonstrates that a boost is essential for development of significant antibody responses in vaccination with FluSure. However, inactivated vaccines with oil-in-water adjuvants have been implicated in the development of vaccine associated enhanced respiratory disease (VAERD) when the vaccine and challenge strain HA are mismatched^26,33^. Therefore, exploring alternative vaccine platforms that protect against heterologous infection without resulting in VAERD is greatly needed in the goal of a universal SIV vaccine. Here we use an Ad5 vector which was previously shown to reduce viral shedding without causing VAERD after challenge with a mismatch SIV strain^34^. In addition, Ad vectored vaccines in swine have shown efficacy in the presence of maternally derived antibodies which limit the efficacy of inactivated vaccines^35–38^. In this study, we demonstrated that the Ad-epigraph swH3 vaccine induces high titers of cross-reactive antibodies after only a single-immunization in swine. In contrast, the Ad-TX98 vaccine induced relatively strain-specific immunity with limited cross-reactivity. Therefore, while an Adenovirus vector can induce strong immunity after only a single immunization, the computational design of the epigraph vaccine contributes to the induction of cross-reactive antibodies.

Vaccination against SIV has two direct benefits: 1) reduction of clinical disease in pigs to prevent economic losses and 2) reduction of viral shedding to decrease transmission within the herd^30^. This reduced viral shedding also reduces the potential for spill-over infections to humans. Our challenge studies in mice have shown that epigraph vaccination, in addition to reducing weight loss, also resulted in the greatest reduction in lung viral titers after challenge with three highly diverse swH3 viruses as compared to the other vaccine groups. This reduction in viral titer could lead to reduced viral shedding and, thereby, reduced intra- and inter-species transmission. However, challenge studies in swine are needed to further demonstrate this efficacy.

The transmission of influenza virus from human-to-swine contributes to the viral diversity of SIV^28,39^. Here, we have shown that epigraph vaccination leads to cross-reactive antibody titers to multiple human H3 isolates after vaccination in both mice and pigs. Importantly, a single immunization of epigraph in swine resulted in cross-reactive antibody titers to 3 of the 7 huH3 isolates while a single immunization with TX98 or FluSure only induced antibody titers to 1 or 0 of the 7 huH3 isolates, respectively. In addition, the epigraph vaccine induced cross-reactive T-cell responses in mice and completely protected mice from weight loss and death after challenge with a lethal huH3 strain. This data indicates that a single immunization with the swH3 epigraph could protect pigs from several strains of human H3, however challenge studies in swine are needed to confirm this.

In addition to strong cross-reactive antibody responses, the Ad-epigraph vaccine induced strong broadly cross-reactive T cell immunity against both swine and human influenza viruses. In mice, we detected an immunodominant T cell epitope, predicted to be a cytotoxic T lymphocyte CD8 epitope, along with multiple subdominant T cell responses, likely CD4 T helper (Th) cells. Additionally, epigraph vaccination of swine induced a strong total T-cell response against four divergent swH3 strains. In humans, cross-reactive T-cells and the development of memory T cells have been associated with long-lasting immunity against influenza virus^40–43^. In contrast, the role of T cells during influenza infection in swine has not been as well defined. Previous studies in swine have demonstrated cross-protection against divergent SIV strains in the absence of detectable antibodies^44–47^. This suggests a role for cross-reactive T cell responses in the protection against SIV which could also result in long-term durable immunity. However, longevity studies in pigs will be required to establish the durability of the epigraph vaccine-induced immunity.

Here, we have demonstrated the efficacy of the Epigraph vaccine designer tool in the development of a broadly cross-reactive swine H3 influenza vaccine. This is the first report evaluating the epigraph algorithm for the design of a universal influenza vaccine. With our promising results, the epigraph design could be applied to human influenza strains to create a human universal influenza vaccine candidate. Our results not only provide a promising swine vaccine, but also a model for human influenza. Indeed, influenza infection in swine share many similarities with influenza infection in humans, such as similarities in clinical symptoms, distribution of viral receptors in the respiratory tract, and subtypes of influenza causing endemic infections^48^. Therefore, swine make an excellent model for the development and testing of universal influenza vaccines. In this study, we demonstrated broadly cross-reactive humoral and cellular immunity after vaccination with epigraph in both murine and swine models. These data support the development of an epigraph vaccine as a universal swH3 vaccine capable of providing cross-protection against highly divergent strains of swH3.

## Supporting information

Supplemental Figures swH3 vaccine

## Acknowledgements

We thank the Biodefense and Emerging Infectious Disease (BEI) Repository, the USDA Swine Surveillance Influenza A virus isolates repository for reagents used in this study. This research was supported by the National Institutes of Health under Ruth L. Kirschstein National Research Service Award 1 T32 AI125207.

## Conflicts of Interest

Eric A Weaver is an inventor and has a patent application submitted and in progress for the Epigraph immunogens used in this study. (Application: 62/734,791, International Application: PCT/US19/52137)

## Notes

### Competing Interest Statement

The authors have declared no competing interest.

### Summary of Updates

The panels a in figure 7 had been inverted. Figure 7 was replaced to accurately represent the data in panels A and B.

