## Supplemental Figures swH3 vaccine for "Epigraph Hemagglutinin Vaccine Induces Broad Cross-reactive Immunity Against Swine H3 Influenza Virus"

| Vaccine | vp/ml | IFU/mL | vp:IFU ratio |
| --- | --- | --- | --- |
| Epigraph 1 | 4.8e11 | 6.0e9 | 79:1 |
| Epigraph 2 | 1.2e12 | 1.3e10 | 90:1 |
| Epigraph 3 | 1.5e12 | 1.3e10 | 117:1 |
| TX98 | 3.6e12 | 5.8e10 | 62:1 |

**Supp. Table 1**. **Infectivity of recombinant Ad viral stocks.** All three epigraph immunogens and the Texas/1998 (TX98) wild type HA comparator were cloned into a replication-defective Adenovirus type 5 (Ad) vector and the virus particles (vp) to infectious unit (IFU) ratio was determined to confirm equivalent stock infectivity.

| Virus Strain | % amino acid identity to | | | | | |
|---|---|---|---|---|---|---|
| | | | | | FluSure Strains | |
| | Epigraph1 | Epigraph2 | Epigraph3 | TX98 | Minnesota /2012 | North Carolina/2012 |
| A/swine/North Carolina/A01270394/2012 (IV-A; FluSure) | 98.6 | 88.7 | 84.3 | 90.1 | 95.4 | 100.0 |
| A/swine/Minnesota/A01270872/2012 (IV-B; FluSure) | 96.8 | 87.5 | 85.7 | 91.0 | 100.0 | 95.4 |
| A/swine/Texas/4199-2/1998 (I) | 91.5 | 87.5 | 84.5 | 100.0 | 91.0 | 90.1 |
| A/swine/Ohio/09SW73E/2009 (IV) | 96.6 | 87.6 | 84.6 | 91.3 | 95.8 | 95.2 |
| A/swine/Manitoba/00446/2005 (IV) | 98.2 | 88.2 | 85.3 | 92.2 | 97.5 | 96.8 |
| A/swine/Ohio/11SW87/2011 (IV-A) | 98.2 | 88.2 | 84.3 | 89.8 | 95.1 | 99.3 |
| A/swine/Wyoming/A01444562/2013 (IV-A) | 97.9 | 88.5 | 84.5 | 89.8 | 95.1 | 98.9 |
| A/swine/Minnesota/A01432544/2013 (IV-B) | 95.2 | 87.6 | 85.9 | 89.1 | 95.2 | 93.8 |
| A/swine/Indiana/A01202866/2011 (IV-C) | 97.0 | 86.7 | 85.2 | 91.0 | 95.8 | 95.6 |
| A/swine/Kansas/11-110529/2011 (IV-F) | 95.6 | 86.6 | 84.6 | 90.8 | 93.8 | 94.5 |
| A/Swine/Colorado/23619/99 (II) | 91.5 | 89.2 | 86.4 | 93.0 | 91.7 | 90.3 |
| A/swine/Colorado/1/1977 (Human-like) | 88.3 | 85.7 | 85.2 | 90.8 | 87.6 | 87.6 |
| A/swine/Texas/A01785781/2018 (Human-like) | 88.2 | 90.8 | 86.4 | 88.9 | 87.8 | 86.7 |

**Supp. Table 2. Percent identity between the vaccine immunogens and the panel of 12 swH3 representative viruses.** The percent identity was determined for each vaccine HA and the 12 swH3 virus strains used the HI assay.

A

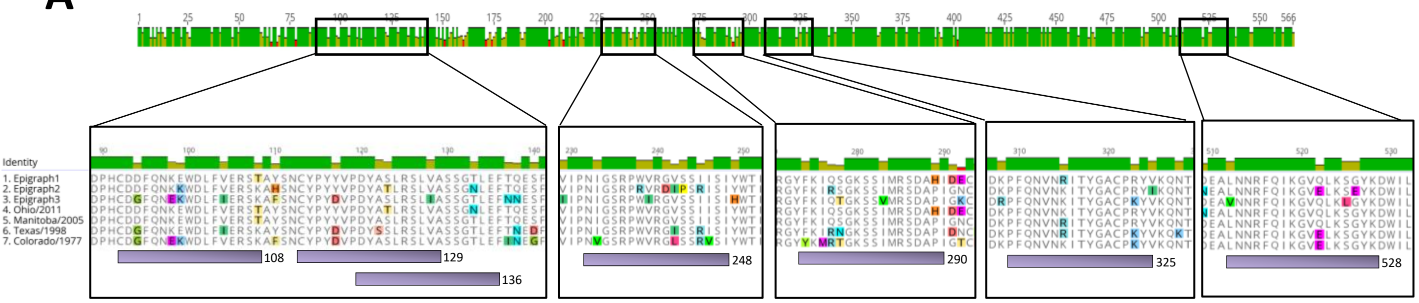

B

| Epitope sequence (120-128) | Virus Strain w/ epitope | Percent of protein sequence matches at 100% identity |
|---|---|---|
| DYASLRSLV | Epigraph 1 | 44.01% (687/1561) |
| | Manitoba/2005 | |
| | Colorado/1977 | |
| DYATLRSLV | Epigraph 2 | 21.20% (331/1561) |
| | Ohio/2011 | |
| DYASLRSLI | Epigraph 3 | 4.16% (65/1561) |
| DYSSLRSLV | Texas/1998 | 3.20% (50/1561) |

**Supp. Fig. 1. T-cell mapping epitope characterization for the swH3 strains.** (A) The protein alignment of each positive peptide (>50 spot forming cells per million) and it's relative location on the HA protein is shown. The peptide number designates the position of the last amino acid in the peptide on the total HA protein. (B) The epitope conservancy of each immunodominant epitope 9-mer (amino acid 120-128) in the swH3 sequence population (1561 sequences) was determined using the Immune Epitope Database and Analysis Resource (IEDB).

| Virus Strain | % amino acid identity to | | | | | |
|---|---|---|---|---|---|---|
| | | | | | FluSure Strains | |
| | Epigraph1 | Epigraph2 | Epigraph3 | TX98 | Minnesota /2012 | North Carolina/2012 |
| A/Aichi/2/68 | 86.7 | 83.9 | 84.1 | 89.0 | 85.7 | 86.0 |
| A/Texas/1/1977 | 89.8 | 86.4 | 86.2 | 92.4 | 89.8 | 89.0 |
| A/Mississippi/1/85 | 91.5 | 87.6 | 86.0 | 94.2 | 90.8 | 90.5 |
| A/Sydney/5/1997 | 92.9 | 88.9 | 86.4 | 94.9 | 92.9 | 91.5 |
| A/Nanchang/933/1995 | 93.6 | 89.6 | 85.3 | 97.0 | 93.3 | 92.2 |
| A/Beijing/352/1989 | 92.0 | 87.8 | 84.8 | 95.9 | 91.7 | 90.6 |
| A/Ann_Arbor/3/93 | 90.0 | 86.3 | 82.4 | 94.5 | 89.7 | 88.8 |

**Supp. Table 3. Percent identity between the vaccine immunogens and the panel of 12 huH3 representative viruses.** The percent identity was determined for each vaccine HA and the 7 huH3 virus strains used the HI assay.

**A**

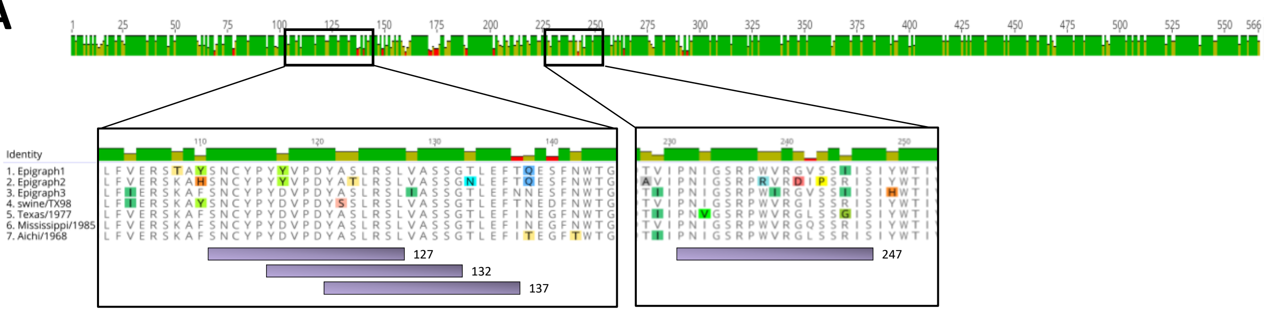


**B**

| Epitope sequence (120-128) | Virus Strain w/ epitope | Percent of protein sequence matches at 100% identity |
|---|---|---|
| DYASLRSLV | Epigragh 1<br>Texas/1977<br>Mississippi/1985<br>Aichi/1968 | 93.68% (5348/5709) |
| DYATLRSLV | Epigraph 2 | 0.16% (9/5709) |
| DYASLRSLI | Epigraph 3 | 1.68% (96/5709) |
| DYSSLRSLV | swine/TX98 | 0.04% (2/5709) |

**Supp. Fig. 2. T-cell mapping epitope characterization for the human H3 strains.** (A) The protein alignment of each positive peptide (>50 spot forming cells per million) and it's relative location on the HA protein is shown. The peptide number designates the position of the last amino acid in the peptide on the total HA protein. (B) The epitope conservancy of each immunodominant epitope 9-mer (amino acid 120-128) in the huH3 sequence population (5709 sequences) was determined using the Immune Epitope Database and Analysis Resource (IEDB).

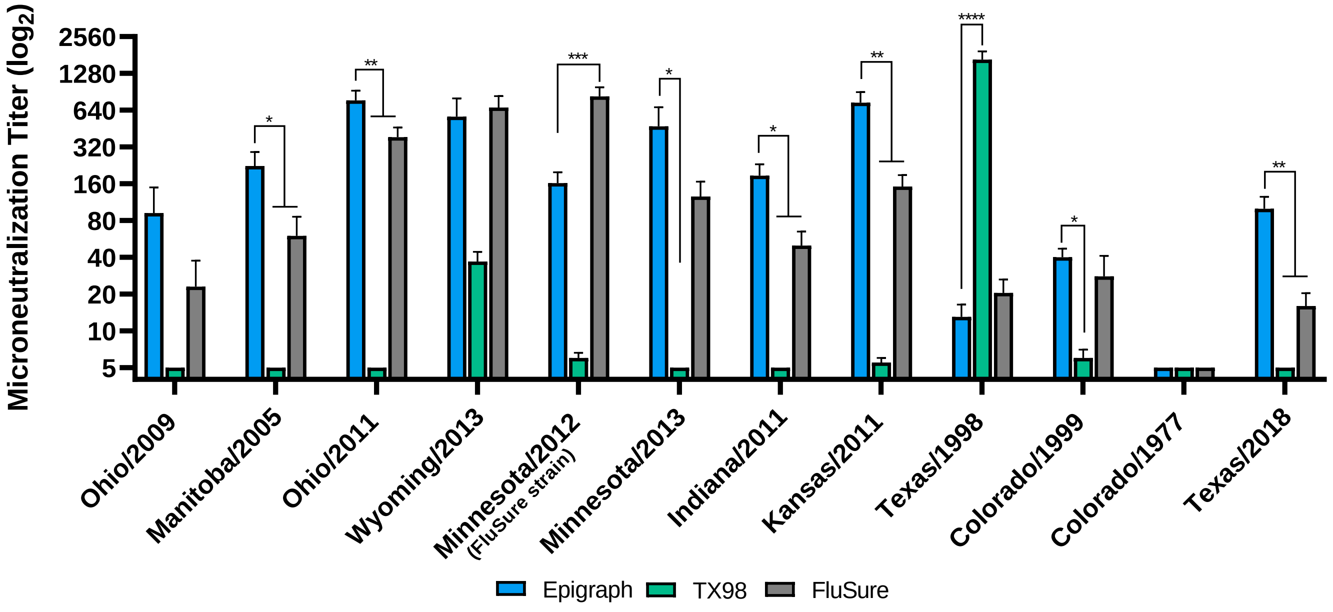


**Supp. Fig. 4. Microneutralization titers after prime boost immunizations in swine.** The swine sera after boosting was analyzed for the ability to neutralize the 12 swH3 strains using a microneutralization assay (one-way ANOVA compared to the epigraph group; *$p < 0.05$; **$p < 0.01$, ***$p < 0.001$).
